# Classification of human physical activity based on the raw accelerometry data via spherical coordinate transformation

**DOI:** 10.1101/686519

**Authors:** Michał Kos, Małgorzata Bogdan, Nancy W. Glynn, Jaroslaw Harezlak

## Abstract

Human health is strongly associated with person’s lifestyle and levels of physical activity. Therefore, characterization of daily human activity is an important task. Accelerometers have been used to obtain precise measurements of body acceleration. Wearable accelerometers collect data as a three-dimensional time series with frequencies up to 100Hz. Using such accelerometry signal, we are able to classify different types of physical activity.

In our work, we present a novel procedure for physical activity classification based on the raw accelerometry signal. Our proposal is based on the spherical representation of the data. We classify four activity types: resting, upper body activities (sitting), upper body activities (standing) and lower body activities. The classifier is constructed using decision trees with extracted features consisting of spherical coordinates summary statistics, moving averages of the radius and the angles, radius variance and spherical variance.

The classification accuracy of our method has been tested on data collected on a sample of 47 elderly individuals who performed a series of activities in laboratory settings. The achieved classification accuracy is over 90% when the subject-specific data are used and 84% when the group data are used. Main contributor to the classification accuracy is the angular part of the collected signal, especially spherical variance. To the best of our knowledge, spherical variance has never been previously used in the analysis of the raw accelerometry data. Its major advantage over other angular measures is its invariance to the accelerometer location shifts.

## 1 Introduction

An important task in medical science is obtaining a thorough and objective characterization of a person’s physical activity. The daily level of one’s activity is highly correlated with widely understood human health and is often used in different areas of medicine as a health and physical fitness indicator. A good example is the problem of comparison of different rehabilitation methods after surgery, where accurate measurements of patients activity level allow to monitor recovery process. In order to ensure objectiveness and high precision of measurements, scientists turned attention to analysis of signal from body-wearable devices such as accelerometers (Bussmann et al. (2001); Atienza and King (2005); Sirard et al. (2005); Boyle et al. (2006); Grant et al. (2008); Troiano et al. (2008); Kozey-Keadle et al. (2011); Choi et al. (2011); Schrack et al. (2014)). Accelerometers measure a three dimensional acceleration vector (*a* = (*x, y, z*)*^T^*) generated by a part of the body to which a device is attached (eg. a hip or a wrist). The measurements are taken with high frequency in the range of 10 to 100 Hz. From the statistical perspective, the signal can be thought of as a nonstationary time series without any explicit pattern with the exception of simple activities such as resting, where the signal is for most part flat, or walking, where the signal exhibits periodicity. There are currently two major sets of methods to analyze accelerometry data. The first set uses the aggregated signal to provide different measures of the energy expenditure, physical activity volume or its intensity (see: Bai et al. (2016); van Hees et al. (2013); Bai et al. (2014)). The second set, which our work expands on, provides classification techniques for the human activity modes (see: Pober et al. (2006); Mannini et al. (2013); Krause et al. (2003); Staudenmayer et al. (2009); Trost et al. (2012); Zhang et al. (2012); Xiao et al. (2015); Urbanek et al. (2015); Straczkiewicz et al. (2016)).

In our work, we develop a novel classification method of human physical activity based on the spherical representation of the raw accelerometry signal. Our proposed methodology categorizes human activity into one of four classes: resting, upper body activities performed in a sitting position, upper body activities performed in a standing position and lower body activities. The method enables classification on the order of seconds. As a result, it recognizes short term bouts such as walking few steps or getting up from a chair. Analysis of a raw signal in the spherical coordinate system is a natural extension of methods which use only the information on the radius. Our approach additionally enables exploration of angular changes of the raw accelerometry data. Naturally, the angular coordinates (*ϕ* and *θ*) depend on rotations of the accelerometer which is illustrated by plots in the third row in Figure 1.1. However, on their basis one can construct rotationally invariant variable called spherical variance 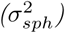. This new rotationally invariant variable measures dispersion of points on a sphere. Furthermore, the spherical variance together with the variance of the radius 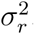 (see fourth row in Figure 1.1) are the most important statistics in the classification process. Many researchers neglect the information contained in the angular part of the signal because of its rotational instability. From that point of view, the spherical variance which is insusceptible to device rotations and calculated by using only angular coordinates, provides additional information on top of the acceleration vector signal. It is important to emphasize that to our knowledge this is a new use of the spherical variance in the context of accelerometry data. In the classification process, in addition to the both types of variance, we have used means of spherical coordinates (*µ*_*r*_, *µ*_*ϕ*_ and *µ*_*θ*_). The mean of the radius (*µ*_*r*_) is the next rotationally independent variable. On the other hand angles means (*µ*_*ϕ*_ and *µ*_*θ*_) enable exploration of accelerometer’s spherical arrangement during performance of different activities. They turn out to be useful particularly when performing within subject classification. We used the summary statistics based on the spherical coordinates to construct classifiers via the decision tree method. We obtained high predictive abilities of different types of human activities. The value of classification accuracy for the strongest models was 90% on the within-subject level and 84% for between-subject case. In order to establish the importance of different variables we used the random forest method.

**Fig. 1.1.**
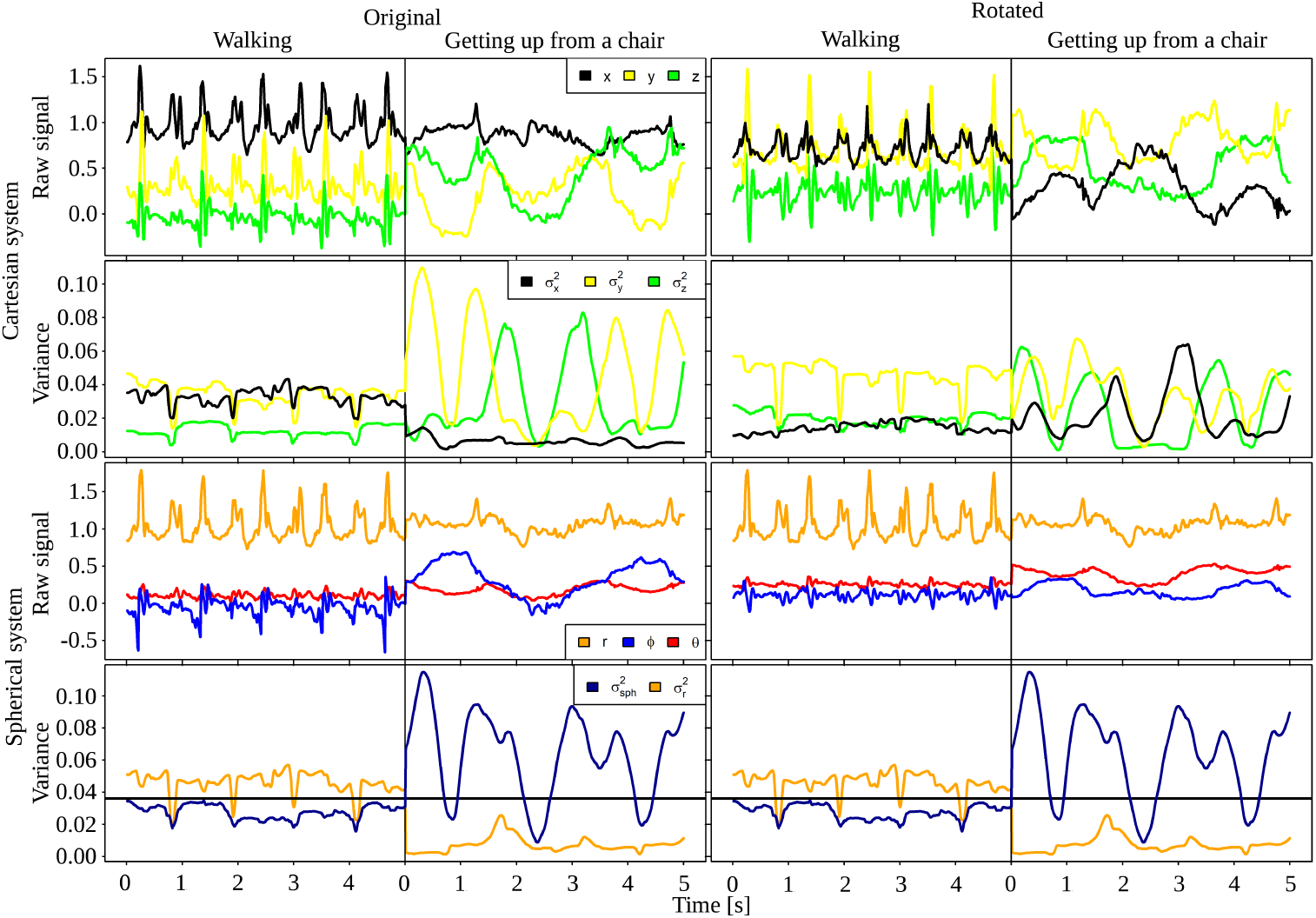
Comparison of signal for walking and getting up from a chair. Two top rows present signal in Cartesian coordinate system. Two bottom rows show the same information in spherical coordinate system. First and third row present raw accelerometry data. In second and fourth row we have variances of coordinates obtained for each time point. For spherical system we present sph spherical variance 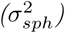 instead of variances for angles. This is one of the most important variables in classification process. In first column (original) we present data in unmodified system. In second column (rotated) we present the same signal which would be obtained if accelerometer was rotated by 45 ^°^ with respect to x and y axes. One can notice that Cartesian coordinate system is sensitive to rotational changes of accelerometer. In case of spherical system both types of variance and radius(r) are rotationally invariant and we can establish thresholds (e.g. black line for variances)

We are not the first to address the issue of identifying this type of human activity. Among earlier works, Pober et al. (2006) proposed two algorithms: the first based on quadratic discriminant analysis and the second utilizing hidden Markov models. Both methods were used to classify four different activities (walking, walking uphill, vacuuming, working at a computer) performed by six participants. Mannini et al. (2013) used support vector machine technique to develop an algorithm that classifies four activity groups: sedentary (resting and low intensity activities), cycling, ambulation (different types of walking) and other activities. Krause et al. (2003) constructed classifiers by utilizing unsupervised clustering and Markov models. Artificial neural networks were used by Staudenmayer et al. (2009) and Trost et al. (2012). Staudenmayer et al. (2009) worked with signals generated by adults between the ages of 21 and 69. In the classification process four activity types were differentiated: vigorous sports, low level activities, locomotion and household activities/other. On the other hand Trost et al. (2012) evaluated young participants between the ages of 5 and 15. They categorized activity into one of five groups: sedentary, walking, running, light intensity household activities or games, and moderate-to-vigorous intensity games or sports. Zhang et al. (2012) proposed an algorithm using combined methods that classified activity as walking, running, household, or sedentary activities.Xiao et al. (2015) introduced technique based on an empirical basis called movelets designed specifically for the analysis of accelerometry data. The procedure classified 5 activity types: standing, lying, walking, upper body activities and getting up from a chair. Urbanek et al. (2015) and Straczkiewicz et al. (2016) proposed methods based on short-time Fourier transformation to detect sustained harmonic walking and the identification of car driving periods in human activity, respectively. Each of the methods mentioned above have different limitations. A number of them classify human activity for intervals longer then 10 seconds (Staudenmayer et al. (2009); Trost et al. (2012); Zhang et al. (2012); Krause et al. (2003)). Consequently, they are unable to recognize short term activity changes which can overlook first signs of a patients recovery such as walking few steps around the house. These are important activities for comparative analysis of different methods of rehabilitation. Other techniques (Zhang et al. (2012); Mannini et al. (2013); Urbanek et al. (2015); Straczkiewicz et al. (2016)) transform data into the frequency domain and use the dominant frequencies for classification. Such features, in order to be useful, require systematic and periodic repetition of activity which is not common in everyday life. An example where such approach is beneficial one can find in Straczkiewicz et al. (2016); Urbanek et al. (2015). In the first paper the authors utilize periodicity in signal which is generated by car vibrations. The second concentrates on harmonic walking which lasts at least 10 seconds. This method enables detection of walking to a bus stop or a shop. However it has problems with classification of walking a few steps around house which can be inharmonic and last less than 10 second. Another major limitation of the aforementioned methods is that they do not use the raw accelerometery data efficiently. Many of the classification methods are based on the extracted feature called “activity count” (Pober et al. (2006); Krause et al. (2003); Staudenmayer et al. (2009); Trost et al. (2012)). This feature is defined by the device manufactures based on a proprietary algorithm. Other published work, which uses the raw accelerometry data, often works exclusively with the statistics constructed from the acceleration vector magnitude (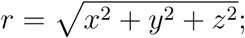 radius) and neglects the angular information (Zhang et al. (2012); Mannini et al. (2013); Urbanek et al. (2015); Straczkiewicz et al. (2016)). These approaches rely on the rotational independence of the activity counts and the vector length, which enables the signal comparisons for different devices and participants. A method which overcomes the aforementioned disadvantages is the movelets technique developed by Xiao et al. (2015). This procedure detects short term activities such as walking a few steps around the house or getting up from a chair. It also incorporates information from the whole signal in the classification process. However, the movelets method uses the Cartesian coordinate system that is sensitive to device’s rotational changes. An illustration of this phenomenon is shown in the top four plots in Figure 1.1. Both columns present the same signal in two Cartesian coordinate systems. However the second show signal which would be measured if the accelerometer was rotated by 45 ^*°*^ around the x and y axes. It is easy to notice that the comparison of both signals would be hard due to horizontal shifts and rescaling of the raw signal. In order to resolve this issue Xiao et al. (2015) proposed a method of data normalization for different participants. However it demands additional information about participants standing and lying periods.

The remainder of our paper is organized as follows. In Section 2, we describe the data set used to illustrate our method and present procedures for the methodology. In Section 3, we summarize the classification performance of our method. The Section 4 contains the discussion, conclusions and future directions of this work.

## 2 Data and Statistical Methodology

This Section covers both the theoretical and practical foundations of the proposed method. We start with the description of the accelerometry data set from the Developmental Epidemiologic Cohort Study (DECOS) used to illustrate our method (Section 2.1). Next, we introduce the accelerometry data transformation into the spherical coordinate system. We also discuss the most appropriate choice of a system for our purpose (Section 2.2). In Section 2.3, we present the definitions and important properties of the variables used to construct classification models. Then we describe the method used to classify (predict) different types of human physical activity (Section 2.4). In Section 2.5 we present construction of the training and test sets. Lastly, we describe the methodology used to compare the constructed models (Section 2.6).

### 2.1 The DECOS data

The predictive abilities of our method have been tested on data collected during the part of the laboratory portion of DECOS (see: Lange-Maia et al. (2015)). In this part of data collection, study participants wore simultaneously three accelerometers placed on the right hip and both wrists. The cohort consisted of 47 older adults (25 males, 22 females) age 70 and older. The participants performed a series of tasks similar to everyday duties. The activities were grouped based on the muscle involvement level of different body parts:

- Resting activities: lying still, standing still, sitting still,
- Upper body activities performed in a sitting position: writing, dealing cards,
- Upper body activities performed in a standing position: washing dishes, dough kneading, folding towels, dressing, vacuuming, shopping,
- Lower body activities: chair stand, fast walk, normal walk, walking on a treadmill 1.5 mph,

Most of the activities were performed for at least 3 minutes. The exception was chair stand which was performed 5 times in a row (duration 10 - 15 s). The start and completion times for each task were recorded by a trained research assistants.

### 2.2 Selection of the spherical coordinate system

Three-dimensional vectors are frequently represented in the spherical coordinate system. To specify such coordinate system, one has to select two mutually orthogonal directions - the zenith and the azimuth. In Figure 2.2 we present the method of choosing the spherical coordinate system, which are applied to all accelerometers. For each participant the zenith and the azimuth corresponded to the natural left and up directions (from participants view), respectively. A vector *P* in this system is represented by three numbers: *r* - radius (signal amplitude), *θ* - called inclination (polar angle), *ϕ* - called azimuthal angle. The choice of the zenith and azimuth directions is arbitrary. However for our selection of coordinate system, the angle *ϕ* for the hip signal has a physical interpretation since it measures the angle of the torso in the plane spanned by the forward and up directions. In order to estimate the zenith and azimuth directions we used few seconds of two activities: standing still and laying still. We assumed that the mean vector for standing still coincide with participants “down” direction and that the mean vector for laying still is close to “backward” direction. A precise description of the spherical coordinate system selection for the collected raw accelerometry data is included in an appendix (see: 5.1).

**Fig. 2.2.**
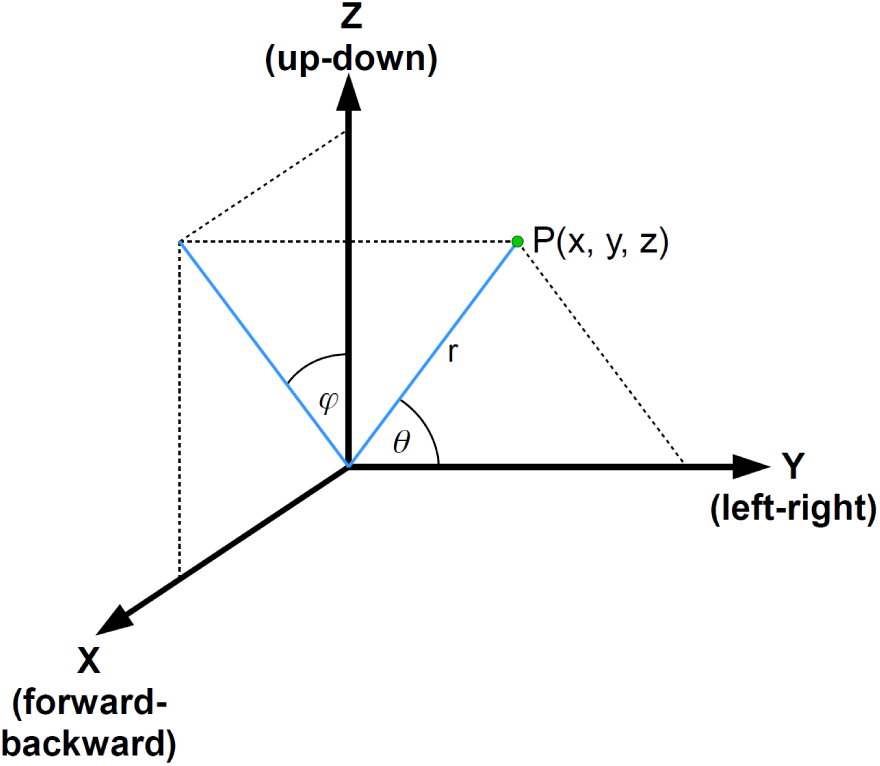
Choice of spherical coordinate system

**Fig. 2.3.**
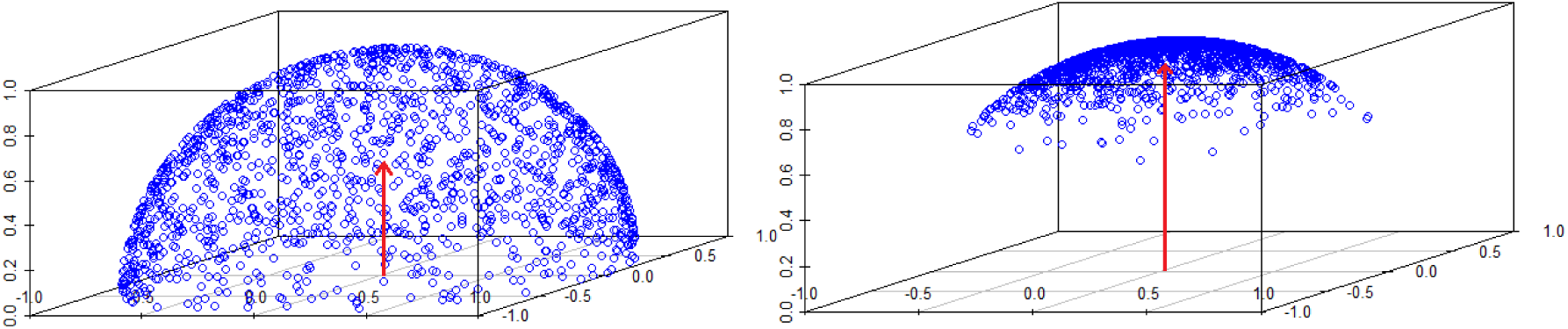
Relation between mean resultant length (R; red arrows length) and points concentration on upper hemisphere. R = 0.92 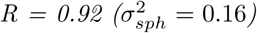 for highly concentrated points (right panel) and has lower value R = 0.5 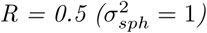 for points spread uniformly on a hemisphere (left panel). For points spread uniformly on a sphere R would be close to 0 (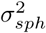 close to maximal value 2).

### 2.3 Description of the candidate classification variables

Accelerometers are typically programmed to collect from 10 to 100 measurements per second (10-100 Hz). Such frequencies exceed the range of frequencies of most human activities. Because of this feature and a fact that the accelerometer raw signal has a high complexity level (nonstationary time series without any explicit pattern) we decided to work with statistics such as the mean value and the variance of spherical coordinates over one-second intervals. More specifically, for each point the statistics were calculated by using 39 preceding points, the point and 40 following points (80 consecutive points = 1 second of signal). This way we obtained new time series with means and variances for different spherical coordinates, which were used to construct classification models. In Table 1 we present variables, which are further used to construct the classification models. The statistics based on radius, mean and variance, are defined in a classical way:

**Table 1:** The description of derived variables used in data analysis

| Notation | Description |
| --- | --- |
| $\mu_r$ | mean value of the vector of magnitude |
| $\sigma_r^2$ | variance of the vector of magnitude |
| $\mu_\phi$ | mean value of the angle $\phi$ |
| $\mu_\theta$ | mean value of the angle $\theta$ |
| $\sigma_{sph}^2$ | spherical variance |

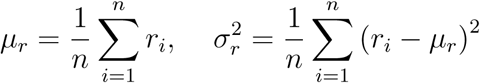

where *n* = 80. The statistic associated with angular information are not that popular and we decided to present theirs definition and derivation of interesting properties. More details about this topic one can find in Mardia and Jupp (2008).

Let us assume that we have a set of points on the unit sphere 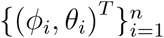. For each point we calculate corresponding Cartesian coordinates *v*_*i*_ = (*x*_*i*_, *y*_*i*_, *z*_*i*_)*^T^* according to classical equations (*r* = 1):

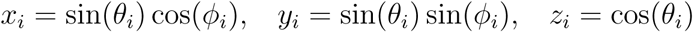

Next we calculate the mean vector:

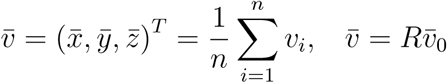

where 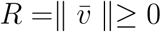 is called the *mean resultant length* and 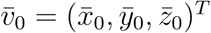 is a unit vector called the *mean direction*. The mean direction allows us to define the mean angles (for details, see: Mardia and Jupp (2008)) according to the following classical equations:

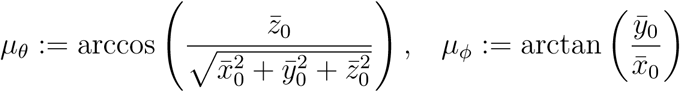

The notion of the *spherical variance* (see: Mardia and Jupp (2008)) is defined in a following way:

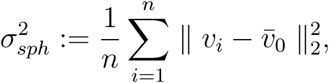

The formula for 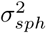 can be transformed into the following relation, from which one can easily determine its value:

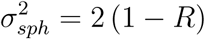

The relation is consistent with intuition. The mean vector of highly concentrated points on a sphere has mean resultant length (*R*)| close to 1 (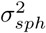 - close to 0; see right panel Fig.: 2.3). On the other hand if points are spread uniformly e.g. on upper hemisphere, then the length of average vector will be smaller and in consequence the spherical variance will have higher value (see left panel Fig.: 2.3). An important property of the spherical variance is a fact that its value does not depend on the coordinate system choice. This feature is very important in comparative analysis. The derivation of above expression for 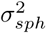 and the proof of its rotational independence are included in an appendix (see: 5.2).

### 2.4 Classification method

In order to investigate classification abilities of derived variables we used the decision tree method. This choice was dictated mainly by two factors. First, the decision tree method enables threshold values determination for the variables separating different groups of activities. This property is important in the interpretation process especially for professionals in medical disciplines. Second, it is relatively easy to explain how the method works for specialist with small experience in statistical field. We present here only a brief summary of a model construction via decision tree method. A full description of a process can be found in Breiman et al. (1984). In a first step the procedure selects one variable that in the best way (in accordance with certain criterion; in our case maximization of gini impurity reduction) separates points from different classes into two subsets. In each consecutive step procedure reiterates the first step (selection of variable and division of a subset) for each “child” subset. The algorithm stops when newly created subset reaches a fixed minimum size or when none of the variables improves class separation at predefined level (e.g. all points in a subset are from the same class). Due to the fact that at each step sets are divided into two subsets a simple and clear method of presenting constructed models are binary trees. In our analysis the decision trees were obtained with *rpart* function from package *rpart* in R environment. During model generation we used the “class” method and default values of parameters which control tree growth. Next we pruned the decision trees by choosing sub-tree with minimal cross-validation error (tree pruning prevents from overfitting of models).

### 2.5 The training and test sets

In our studies, we build a classifier for four basic groups of activities. This was done on two levels - within and between-subject. First, we explored behavior of models constructed on data for individual person (within-subject level). In that part of analysis we have build models for 39 participants separately (8 participants were excluded due to errors in the data e.g. lack of labels of some of the activities). On the other hand, on between-subject level we investigated the decision trees grown on signal from part of the participants and applied to the data collected from the rest of entrants. In that part of study we worked only with signal from right-handed participants (34 persons). Table 2 presents division of activities into groups and the training and test sets sample sizes on within-subject level.

**Table 2:** The activities chosen for model examination

| Group | Activity | Trainig set | Test set |
| --- | --- | --- | --- |
| Resting | laying still,<br>standing still,<br>sitting still | 0.5 minute,<br>(2400 points) | 2.5 min |
| Upper body<br>(siting) | writing,<br>dealing cards | 0.5 min | 1.5 min |
| Upper body<br>(standing) | washing dishes,<br>dough kneading,<br>dressing,<br>folding towels,<br>vacuuming,<br>shopping | 0.5 min | 5.5 min |
| Lower body | chair stand,<br>normal walk,<br>fast walk,<br>walking on a treadmill 1.5mph | 0.5 min | 2.75 min |

In order to construct training and test set we used the middle minute interval (out of three minute long intervals) of each performed activity (with exception of “chair stand”). We worked with middle minute of each activity to capture its routinized and stable part. Moreover, this approach allows to reduce the influence of errors in mislabeling activity start and ending points (this issue appeared especially for tasks performed at the end of experiment). The duration time for “chair stand” activity for different participants was around 10 - 15 s. Because of that we could not treat this activity in above specified way and we used the whole interval for this task. In next step we divided each selected interval into two sub-intervals separated from each other by one second of signal. The training set was a union of first sub-intervals for different activities and has been selected in a way that provides equinumerous group representation - 30 seconds on each activity group. In consequence, the equinumerosity condition determined the length of sub-intervals associated with training set, which was 30*s/k*, where *k* was the number of activities in the group. The collection of remaining sub-intervals determined the test set. Naturally, the construction of test set caused unequal activity groups representation. This problem was taken into account in prediction process (for details see below).

Taking into consideration that on one hand we would like to classify short term action and on the other our signal resolution is 80 Hz, we decided to predict the labels of the activity groups with an accuracy of one second. The exact process of prediction had the following form. First, on the basis of constructed variables 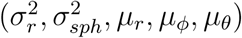 we build the decision tree on a training set. In second step, we divided test set into disjoint 1 second long intervals. Next for each point within interval via constructed decision tree we predicted the label of an activity group. Finally, the predicted label for the entire second was determine as the most recurrent predicted label for all points in interval (via majority voting method). In order to quantify predictive abilities we used classification accuracy notion (the ratio of the number of correctly classified one second intervals to the total number of predictions in percentage scale). The overall classification accuracy was estimated as an average of classification accuracy for all activity groups. This was done due to differences in the sizes of test sets for various activity groups.

On between-subject level the decision tree was constructed by using a set consisting of 4 minutes of signal from each participant. Each minute (out of 4 minutes) corresponded to different activity group and was a union of 60*s/k* long intervals associated with task from a group (k - number of activities in a group) with exception of lower body group. For lower body group the minute consisted out of whole interval for “chair stand” and equally represented remaining activities[(1 min − “chair stand” duration)/(*k* −1)]. The intervals in both situations were chosen so they would be exactly in the middle of activity (equal distance to starting and ending task moments). In order to explore predictive abilities of the method we performed 5-fold cross-validation. This was done in the following way. In the first step we divided randomly participants into 5 groups (4 groups of 7 participants and 1 group of 6 entrants). We calculated the overall classification accuracy for each group separately. This was done by constructing decision tree on data of participants from complementing 4 groups and applied to signal of entrants from specified group. Lastly, we averaged results to calculate estimate of overall classification accuracy. For each group the training set was obtained in the same way - according to above specification (the only difference was that we used here a subset of all participants). On the other hand, the test set was a union over participants from a group of 1 (middle) minutes of activities that were performed for 3 minutes (14 activities) and whole intervals for “chair stand” (14 min per person). The process of classification was analogous to the that for within-subject case.

### 2.6 Comparison of classification models

In our work we addressed a few important issues. One of the most basic is the question about importance of information contained in angular coordinates. Others have worked only with the radius. In this context it is essential to investigate weather our approach provide statistically significant improvement of classification accuracy. The fact that in our experiment participants wore simultaneously three accelerometers (on hip and both wrists) enabled the comparison of classification accuracy of different device placements. Specifically, we compared classification accuracy of models using information from each device separately and trees utilizing information from more than one accelerometer placement. In order to compare mutual significance of accelerometer placements and importance of information contained in angular coordinates, we have worked with 21 different model types. To explore ‘placement level’, we divided models into 7 groups. First 3 were build on the basis of signal from each device separately denoted by H, L and R (respectively: hip, left wrist, right wrist). Next 3 used combined information from two accelerometers (HL, HR, LR) and the 7-th group used signals from both wrists and hip simultaneously (HLR). On ‘angular information level’ we had 3 groups. First, based on information from radius 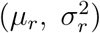 denoted by *’rad’*; second, constructed on invariant variables 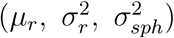 denoted by *’inv’* - strongest invariant model; third, build on all statistics 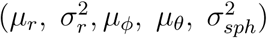 denoted by *’all’* - rotationally dependent. Simultaneous usage of both levels gives us 21 different model types, e.g. ‘*HL inv* ‘ is a model that uses variables 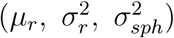 (*’ · inv’*) obtained on a basis of signal from accelerometers placed on a hip and left wrist(*’HL · ‘*).

At within-subject level with each participant we had associated 21 different models. In order to compare mean classification accuracy we fitted following linear mixed model:

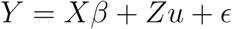

where: *Y*_819×1_ - a known vector of obtained classification accuracy for different models and participants, *X*_819×21_ - design matrix with 0 − 1 entries characterizing method of classification. Each matrix X column correspond to one of 21 methods of constructing decision trees (*X*_*ij*_ = 1 when j-th model was applied). *β* = (*β*_1_, …, *β*_21_)*^T^*- an unknown vector of the classification accuracy means for each method (fixed effects), *Z*_819×39_ - design matrix with 0 − 1 entries describing participant associated with obtained classification accuracy, *u* = (*u*_1_, ⋯, *u*_39_) - an unknown vector characterizing participant influence on the method (random effects), _819×1_ - an un-known vector of random errors. To perform the means multiple comparisons we used Tukey contrast method applied to presented linear mixed model. We used *lmer* function from *lme4* R-package Bates et al. (2015) to fit linear mixed model and *glht* function from *multcomp* R-package Hothorn et al. (2008) (with parameter *linfct* = *mcp*(*method est* = *Tukey*)) for means comparison.

Another addressed issue is the problem of establishing which of the derived variables are responsible for model efficacy. In order to explore it, we used the random forest method, which enabled calculation for each of the so called Mean Decrease Gini. The Mean Decrease Gini is a measure of variable importance. It is calculated as the averaged decrease of Gini impurity (not to be confused with Gini coefficient) in nodes fractured on established predictor across all random forest (for details see Breiman (2001)). The variable importance was calculated on within and between-subject level. On within-subject level for each participant a random forest containing 100 000 trees was grown on the training set. On between-subject level we grown random forest of 50 000 trees. Each out of 34 participants was represented by 24 seconds of signal. The activity groups contained 6 seconds of data from each entrant. This approach provided equinumerous sizes of activity groups. Each activity in a group was represented by 6*/k* seconds long interval, where k is a number of activities in a group. For each activity the intervals left bound coincided with the left bound of the middle minute or with activity starting point (only for chair stand - small duration). In order to construct the random forests we used *randomForest* function with default settings from *randomForest* R-package Liaw and Wiener (2002).

## 3 Results

In this Section, we present the results of the accelerometry data analysis at the within and between-subject level.

### 3.1 Results for models constructed independently on each participant

Section 3.1.1 provides information about classification abilities of models associated with the different accelerometer placements. Furthermore it present results showing significance of angular coordinates. It also shows that on the within-subject level, from the classification perspective it is sufficient to use a model constructed on the whole information from the hip or a model built on rotationally independent statistics from the hip and right wrist. We obtain this result by classification abilities comparison of models constructed on one accelerometer to models that uses information from more than one device. Results presented in Section 3.1.2 addresses an important question of the individual variables significance in classification problem. In Section 3.1.3 we provide information about interesting properties and similarities between decision trees obtained for different participants.

#### 3.1.1 The classification accuracy comparison

In this Section we explore two issues: mutual significance of classification models constructed on the basis of signal from different placement of accelerometer and angular information importance. In the Figure 3.4 on the y-axis we have classification accuracy (the ratio of the number of correctly classified one second intervals to the total number of predictions in percentage scale). Each boxplot contains information about performance of 39 models build independently for different participants. The x-axis informs about the placement of accelerometers (capital letters H, L, R - respectively hip, right and left wrist) and statistics used to construct the models (’all’ - all variables; ‘inv’ - variables rotationally independent 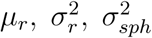 – variables constructed on the radius 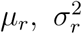). The colors of boxplots are the same for models constructed on information from the same accelerometer placements.

The picture is divided by two vertical lines into three areas. First area (first 9 boxplots) characterize behavior of models constructed on data from each accelerometer separately (H, L, R). Second area (next 9 boxplots) give information about trees build on signal from 2 devices (HL, HR, LR). Last 3 boxplots uses data from all three accelerometers simultaneously (HLR). Let us concentrate at the beginning on the first area that is of particular interest due to a fact that in most applications people use only one accelerometer. If we take a closer look we can notice a few important properties. First of all a comparison of models associated with different placements of device and containing whole information (first in each group - ‘all’) show that the strongest classification accuracy (median equals 90%) has model corresponding to a hip. It is followed by decision tree associated with right wrist (median equals 84%), which was a dominant hand of most of participants. The least effective model correspond to a left wrist (median equals 80%). This points to particular importance of information from a hip.

On the other hand, if we compare boxplots in each color group separately, it is easy to notice that angular variables generally improve predictive abilities of models. This is confirmed by collected in the Table 3 results of performed twoside paired t-tests. Analysis shows that for the signal from the hip particular importance have means of angles and that the spherical variance do not improve significantly classification accuracy. For the left wrist the situation is reversed. Spherical variance ameliorates models performance and angles means do not. In case of the right wrist both modifications significantly improves classification accuracy.

**Fig. 3.4.**
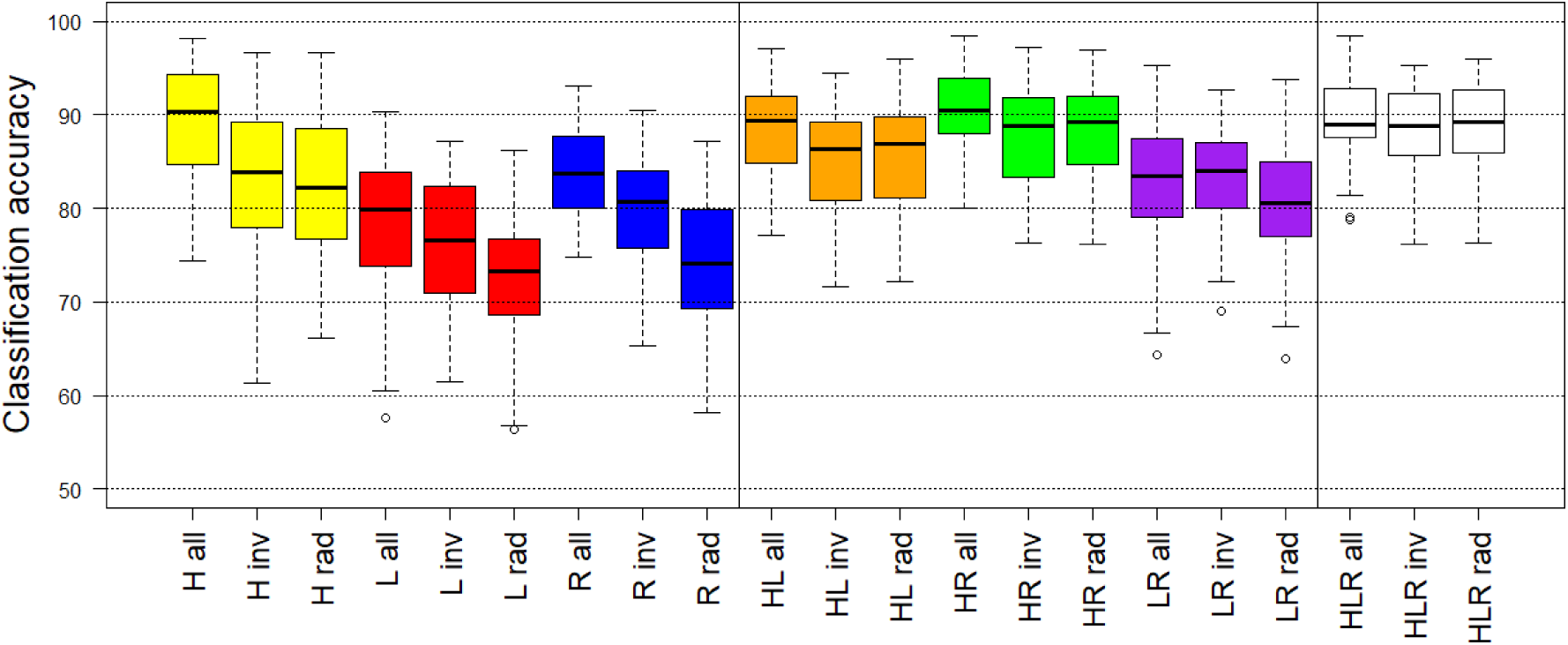
Classification accuracy for 39 participants

**Fig. 3.5.**
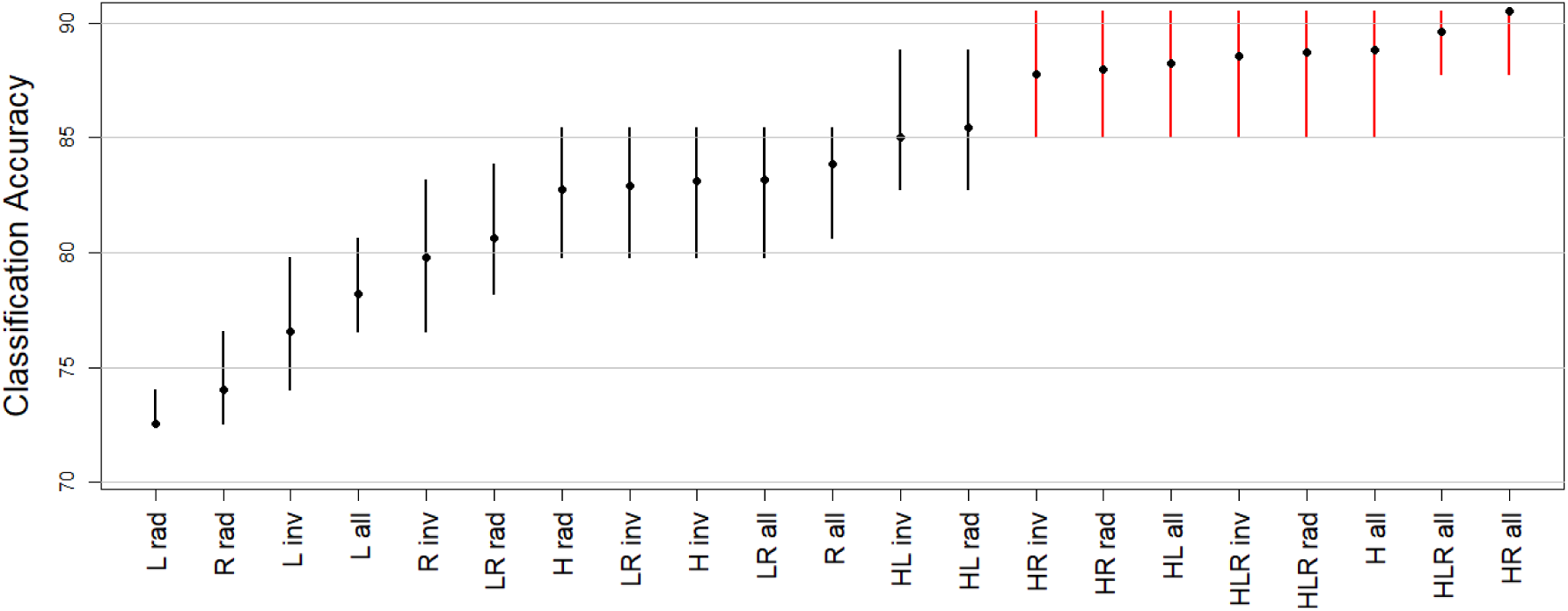
Multiple comparisons of mean via Tukey contrast method. The dots correspond to mean classification accuracy over 39 participants obtained by each model. The whiskers represent the interval in which mean do not differ statistically from means of other models.

**Fig. 3.6.**
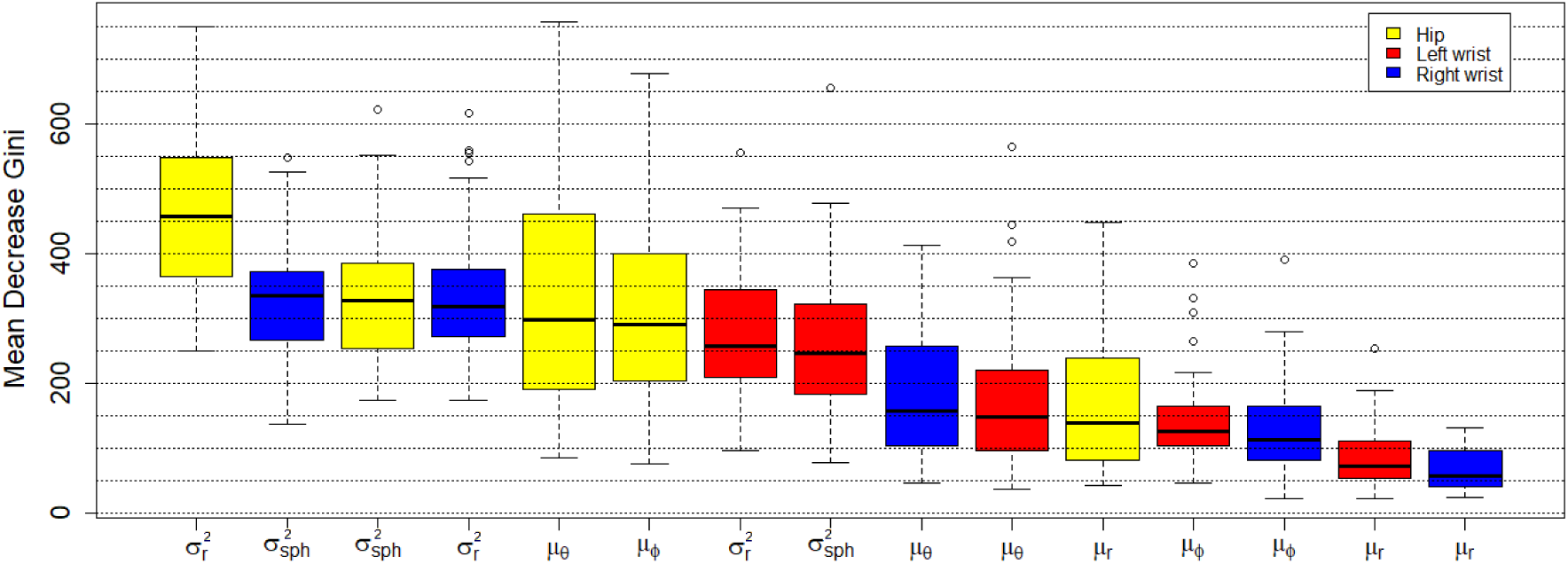
Comparison of variable importance for statistics obtained on the basis of signal from different accelerometer placements.

**Fig. 3.7.**
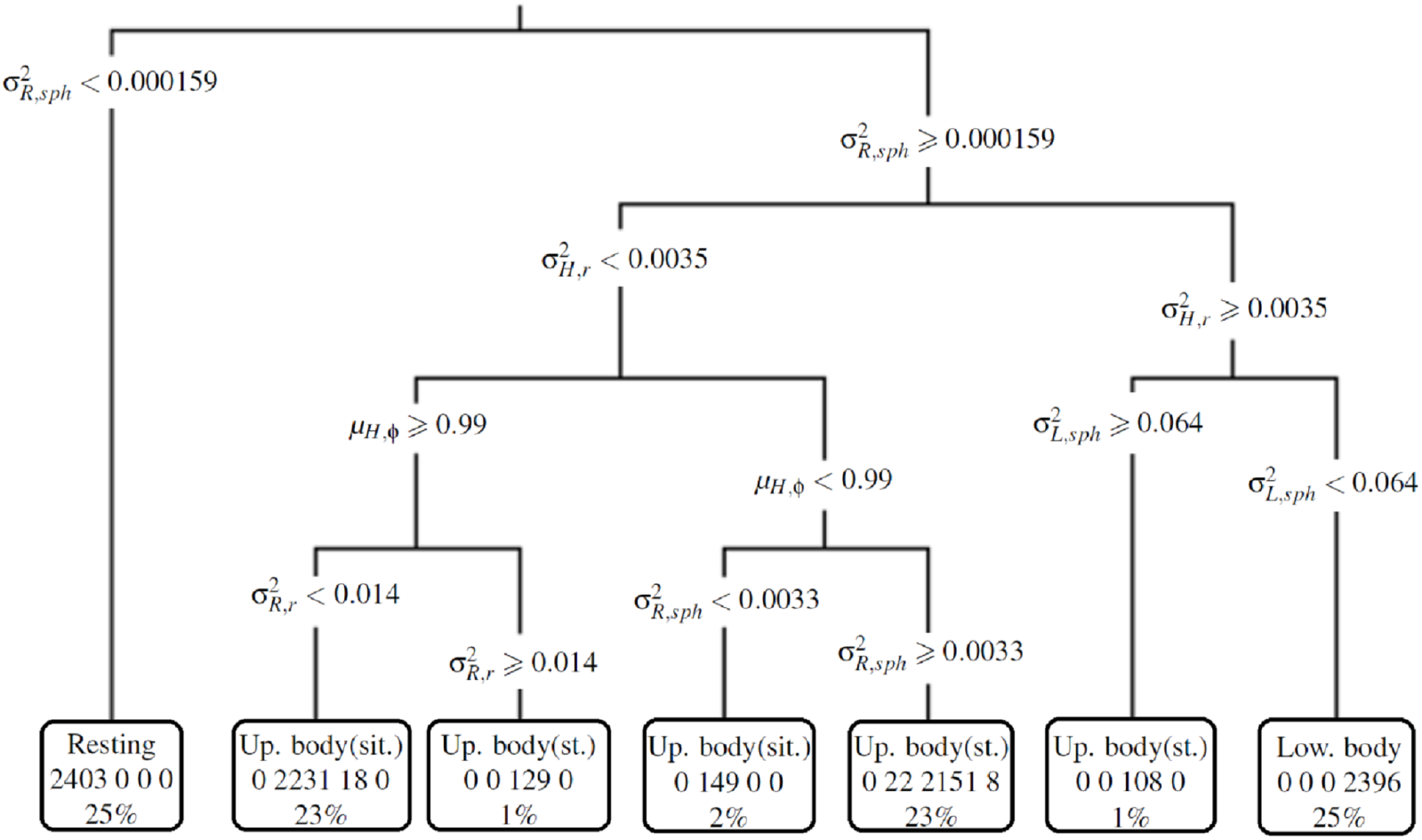
Example of decision tree from one of the participants.

**Fig. 3.8.**
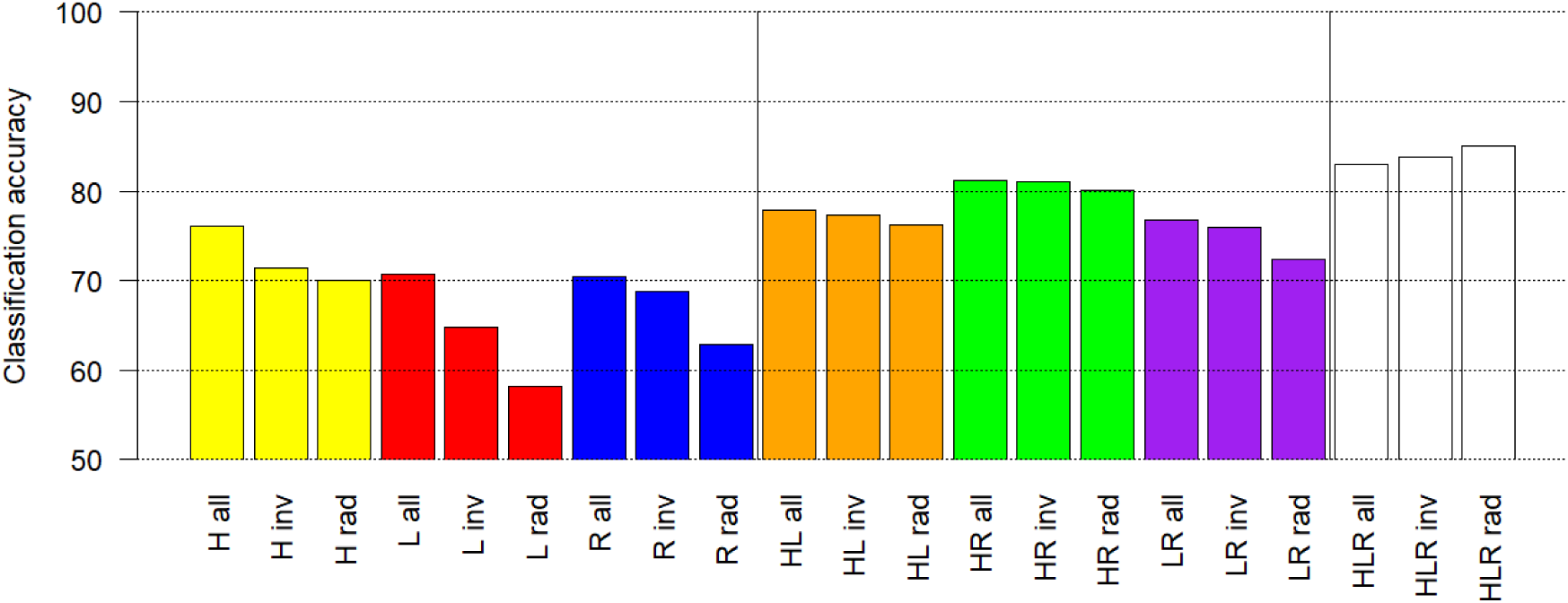
Results of classification accuracy for models on between-subject level.

**Fig. 3.9.**
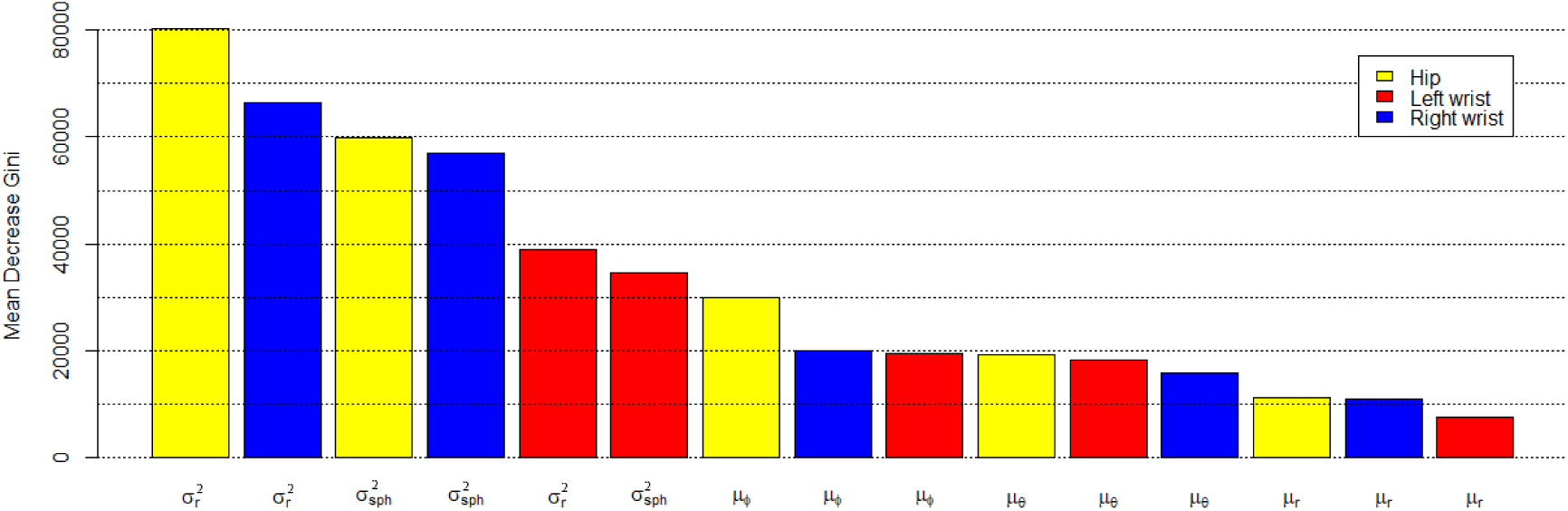
Comparison of variable importance on between-subject level.

**Fig. 3.10.**
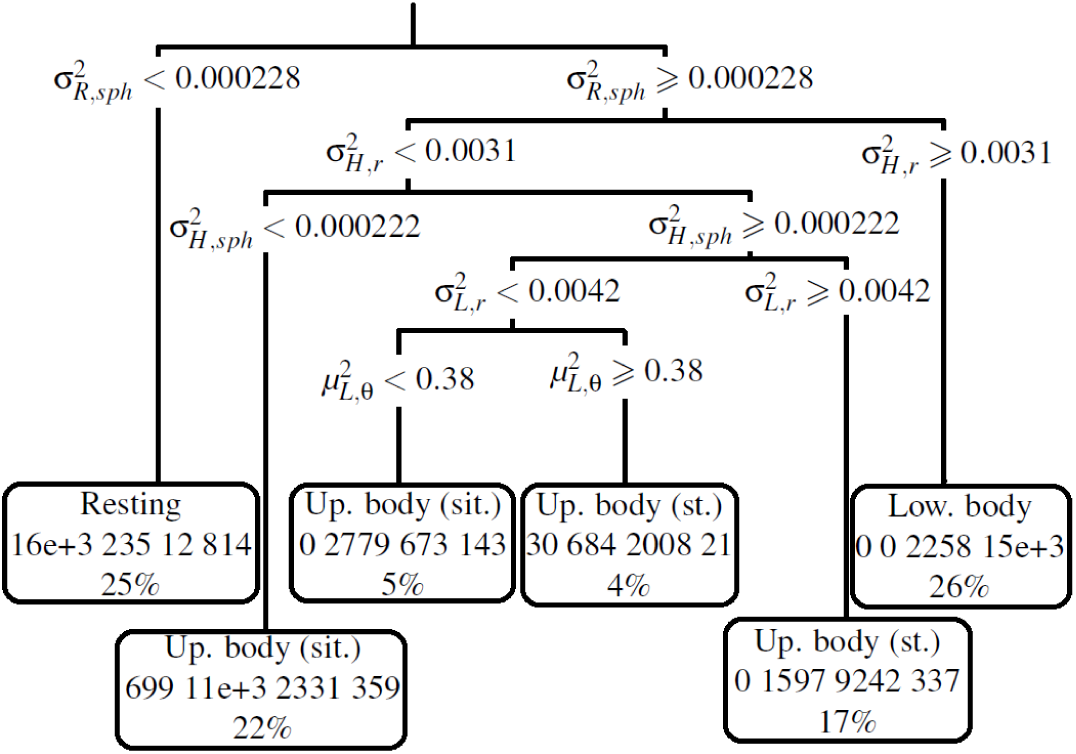
Decision tree containing information from all three accelerometers and constructed on the basis of signal from 34 right-handed participants.

**Table 3:** Results of classification accuracy mean comparison performed for each boxplot color group (paired t-tests).

|  | Mean of the difference | 95% conf. interval | p-value |
| --- | --- | --- | --- |
| H all vs. H inv | 5.71 | (4.13, 7.29) | $8.82 * 10^{-5}$ |
| H inv vs. H rad | 0.35 | (-0.40, 1.09) | 0.35 |
| L all vs. L inv | 1.63 | (-0.55, 3.80) | 0.14 |
| L inv vs. L rad | 4.01 | (2.47, 5.56) | $5.99 * 10^{-6}$ |
| R all vs. R inv | 4.06 | (2.27, 5.86) | $4.80 * 10^{-5}$ |
| R inv vs. R rad | 5.76 | (4.22, 7.27) | $4.03 * 10^{-9}$ |

The above observations suggest that if we have only one accelerometer the best place to attach it is at the hip. On the other hand, many people find it more convenient wearing the device on a wrist (similarly to a watch) rather then on a hip which requires to wear a belt. For such a case it is better to attach the accelerometer to a dominant hand. Let us consider now a situation where we can use more then one device (second and third area). It is visible that the strongest models have similar classification accuracy median to the tree constructed on all variables associated with device from a hip (’H all’). From that perspective we do not gain any value by combining information from more then one device. On the other hand we can see that the IQR of ‘H all’ model is larger in comparison with IQR of strongest models which use information from more then one device. So the classification accuracy is more concentrated around the median for trees combining information from different accelerometers. In case of models built on signal from two accelerometers we can see that the best are ‘HR’ (green) followed by ‘HL’ (orange) and the least effective are ‘LR’ trees (purple). In the case of models containing information from the hip (’HR’, ‘HL’) we can notice that adding spherical variances do not ameliorate classification accuracy (relation between ‘rad’ and ‘inv’). On the other hand the angles means seems to improve classification abilities of models. This behavior is probably a consequence of the dominant role of the signal from the hip for which we had similar conclusions for models constructed on each accelerometers placement separately. In case of models constructed on information from both wrists we observe the reverse situation. Spherical variances improve classification accuracy and angles means do not. For trees build on all three accelerometers placements the differences are insignificant.

In the Figure 3.5 we present results of multiple comparisons of mean via Tukey contrast method. The dots correspond to calculated mean classification accuracy and the whiskers represent the interval in which mean do not differ statistically from means of other models. The models were sorted by the mean. It is easy to identify that the 8 strongest models do not statistically differ. We can observe that the signal from a hip accelerometer is sufficient to obtain the best classification accuracy (’H all’ model). It is visible also that the most parsimonious rotationally invariant model (in group of 8 best models) uses 2 devices and variables constructed on the radius (’HR rad’ model).

#### 3.1.2. The variable importance

The Figure 3.6 shows summary results for 39 participants. On the y-axis we have the values of the Mean Decrease Gini and on the x-axis we have the names of the variables. Each boxplot contains information about values of importance measure for participants. Their colors correspond to the accelerometer placement. The boxplots were ordered by the values of medians. If we look at the first six most important variables we can see dominance of device placed on the hip. It is visible also that the variance of radius associated with the hip is a particularly strong predictor. This shows high importance of signal from a waist. There exist certain hierarchy in each group of boxplots with the same color. The two types of variance are always the strongest variables. Next best are the averaged values of the angles and the weakest predictors are always the means of the radius. Furthermore one can notice that in the group of the first eight strongest predictors, six of them are the different types of variance and their medians are approximately on the same level or higher then the upper quartiles of boxplots for rest of variables. This indicates the high importance of two types of variances. We would like to point out one more observation. If we turn our attention to boxplots corresponding to signal from right wrist (blue color) one can notice that the mean decrease gini for spherical variance is higher than for the variance of the radius. This probably explains relatively high classification accuracy improvement for comparison of models from previous Section. This observation has practical importance. Often in real life accelerometers are worn similarly to a watch. Taking this into account one can see that limitation exclusively to radial part of signal have strong undesirable influence on data analysis.

#### 3.1.3 Decision trees

In this Section we present observations about decision trees constructed on information from three accelerometers. In a natural way models vary for different participants. However to a certain extent they exhibit similar behavior.

In Figure 3.7 one can see a sample decision tree. At each level, points were divided into two subsets with respect to a certain value of one of the variables. At the end of each path we have a leaf informing about the label of activity group, numbers of observations from training set that were assigned to a subset and the percentage of all observations from training set that fall into a subset.

The models for different persons had from 3 to 6 levels. However most of the decision trees had 4 or 5 levels (41.0% and 39%, respectively). In Table 4 one can find information about the frequencies of variables appearance from different placements of the accelerometers. It is again visible dominant role of signal from the hip and certain advantage of the right wrist over the left. Table 5 presents how often different variables appeared. Due to the dominance of signal from the hip we decided to compare frequencies separately for each accelerometer placement. It is noticeable that for the hip three variables with similar appearance frequency dominate. For the right wrist we observed particular importance of the spherical variance followed by medium appearance frequencies of angles means and variance of the radius. For the left wrist dominant role has *µ*_*ϕ*_ followed by 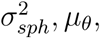, and 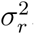. For all three accelerometer placements it is visible low importance of information contained in *µ*_*r*_. An interesting behavior is observed when comparing variances versus means usage on different tree levels.

**Table 4:** Decision trees - placement of accelerometer

| placement | percentage |
| --- | --- |
| hip | 56.1% |
| right wrist | 24.5% |
| left wrist | 19.4% |

**Table 5:** Decision trees - frequencies of variables usage for different accelerometer placement

| variable | hip | right wrist | left wrist |
| --- | --- | --- | --- |
| $\mu_\phi$ | 31.7% | 17.7% | 30.6% |
| $\sigma_r^2$ | 28.2% | 22.6% | 20.4% |
| $\mu_\theta$ | 27.5% | 22.6% | 22.4% |
| $\sigma_{sph}^2$ | 7.7% | 33.9% | 26.5% |
| $\mu_r$ | 4.9% | 3.2% | 0.0% |
| total | 100% | 100% | 100% |

Even though there are more mean variables (60% - means; 40% - variances) we can notice the dominance of variances near the root of the trees (70% on first level). This points again to particular significance of the variances (the decision tree method chooses at each level the strongest variable).

### 3.2 Results for population model

In this Section we present results of classification accuracy on between-subject level. In natural way classification accuracy of model for that case was lower. It is a consequence of a fact that on within-subject level we didn’t have to deal with differences between participants such as e.g. age, height, gender. To make our analysis more immune to entrants individual features we worked only with right-handed participants (34 out of 39 subjects). The Section is organized analogously to the previous one. First, we explore the importance of angular information and compare classification accuracy for models combining information from different accelerometer placements(3.2.1). Next we present results of variable importance (3.2.2) and finally (3.2.3) we present decision tree constructed on signal from all 34 participants.

#### 3.2.1 The classification accuracy comparison

In Figure 3.8, we present the comparison of classification accuracy for models constructed on the basis of signal from different accelerometer placement. The figure is organized in a similar way to the Figure 3.4, which presents the same quantities for within-subject case.

On the y-axis we have classification accuracy and the x-axis contains information about accelerometer placements (capital letters H, L and R for respectively hip, right and left wrist) associated with variables used in model construction (’all’ - all variables; ‘inv’ - variables rotationally independent 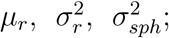; ‘rad’ - variables constructed on the radius 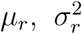). The figure is divided by two vertical lines into three parts. First part shows the results for models constructed on each accelerometer separately. Second part contains information about models utilizing signal from two device placements and the third part displays the results for trees build on all three accelerometers simultaneously. At the beginning let us turn our attention to the first part. Similarly to the within-subject case it is noticeable that signal from a hip has the highest classification accuracy (76%). The strongest models for right and left wrist are on approximately the same level (70%). It is visible also that additional usage of the angular variables increases classification accuracy. The strongest influence is observed for the left wrist where spherical variance increases classification accuracy by 7% and angles’ means provide further improvement of 5.5%. Strong effect is also noticeable for the right wrist. Spherical variance in this case gives 6% increase in classification accuracy. The influence of the angles’ means is also positive however it is rather small (1%). For a hip, effect of angular variables is also small (overall gain 3%).

In the case of the second and third area where we have results for models constructed on information from more than one accelerometer, we can notice that classification accuracies are higher than in the first part. Furthermore, it is visible that the best classification models are obtained when we use all three devices simultaneously (84%). This shows that on between-subject level it is important to use information from different accelerometer placements. On the other hand it is noticeable that significance of angular variables for this case is negligible and that it is sufficient to use only statistics build on the radius. In the case when we have only two accelerometers the best results are obtained for a hip and a right wrist (dominant hand) combination (80 - 82%). There is also noticeable positive influence of the angular variables for the combinations of two devices. The effect is small when models contain information from a hip (’HL’, ‘HR’ - overall 2%) and considerable for models constructed on a left and right wrist (’LR’). For this case the spherical variances increase classification accuracy by 4.5% and the angles means do not give further improvement.

#### 3.2.2 The variable importance

In Figure 3.9 we present results of variable importance analysis for currently considered model. The figure is organized in an analogous way to previous case.

One can notice that figure resembles results from first part of analysis. Again different types of variances have dominant role. The mean values of the radius have the smallest effectiveness and in the middle are the angles means. In consequence the structure within each color group is also preserved. One can notice that if we would look at the same statistics from different accelerometers always the signal from a hip is on first place. This again points to crucial role of signal from a waist.

#### 3.2.3 Decision trees

In Figure 3.10 we present the decision tree containing information from all three accelerometers and constructed on the basis of signal from 34 participants. It is easy to notice the dominance of different variance types (only ones the mean of *θ* was used). This suggest that importance of means has subject-specific character and that it is better to use rotationally independent variables on between-subject level.

On first two levels of tree, model separates totally resting and lower body group from others. The resting activities are isolated by very low values of spherical variance from right wrist. This behavior suggest quiescence of dominant hand and seems to be consistent with intuition (quiescence of dominant hand imply resting nature of activity). On the other hand lower body group is separated from others by high variance of radius from a hip. This is also quite intuitive behavior. High values of variance from a hip suggest vigorous movement of waist (e.g. walking or chair stand). The hardest to separate are upper body activities and as one can notice the structure of decision tree complicates for this two groups.

## 4 Discussion

We proposed and evaluated a novel classification method for different human activity types. Our method is based on a spherical representation of the raw accelerometry signal. The procedure enables accurate classification of short-term activities which is crucial for free-living physical activity assessments. The transformation to a spherical coordinate system enabled introduction of the spherical variance; a summary not used before in the accelerometry data context to the best of our knowledge. Spherical variance is rotationally invariant which is an important feature in comparative analysis of signals from more than one device. Moreover, the results of the analyses indicate high importance of this variable in classification context. Our analyses show that variables characterizing angular changes in accelerometry data significantly improve classification accuracy of the models constructed based only on the information from the radial part of a signal. This property was observed on both within- and between-subject level.

The variable importance analysis (see Figure 3.6 and 3.9) revealed specific hierarchical structure among the extracted features. For both within- and between-subject level as well as each accelerometer placement, the most important features were the variance of the radius and the spherical variance. Next best were the means of the angular coordinates, whereas the radius means had the weakest influence on the classification accuracy. On between-subject level variable importance summarized above is even stronger than for the within-subject level.

The analysis revealed that from the classification accuracy perspective for within-subject level, it is sufficient to use signal from one accelerometer placed on a hip (90%, see Figure 3.4 and 3.5). However, this model uses angle means and in consequence it is important to attach the device each time in exactly the same position, since the angle means are rotationally dependent and hence incomparable for different spatial accelerometer placements. To obtain rotationally independent model with similar classification accuracy we had to use data from two accelerometers placed on a hip and a right wrist. Often, people find it more convenient to wear the device on a wrist (similarly to a watch) rather than on a hip requiring an attachment to a belt. When only one device is worn on a wrist, higher classification accuracy is achieved for the accelerometer placed on a right hand (84%). However, such device placement and modeling strategy uses angle means and hence it is important to pay special attention to the device placement.

For the model at the population level, the best classification accuracy is achieved when data from all three accelerometers is used simultaneously (84%, see Figure 3.8). In this case, the most important predictors were rotationally independent. The best placement combination for two devices is a hip and a right wrist (82%). In this case classification accuracy for full model and rotationally independent predictors is also very close and in consequence it is sufficient to use a model with rotationally independent predictors. When only one device is worn, the best accuracy is obtained for the accelerometer placed on a hip (76%) and next best for the device worn on a dominant hand. However, both models use rotationally dependent predictors, therefore the accelerometer should be attached each time in exactly the same way.

An analysis of decision trees constructed using data from all three accelerometers for within-subject case show certain similarities among them. Optimal trees have very few branches, with majority of them having 4 or 5 levels (80%). Most often the variables extracted from the hip-worn accelerometer are chosen (56%, see Table 4). This again points to a particular importance of the signal from a hip. In a group of variables associated with the device attached to a hip we observed that variables chosen most often were variance of radius and two angle means (see Table 5). In the case of a right wrist the spherical variance was dominant and other important variables included variance of radius and two angle means. For the left wrist, the importance of angle means and spherical variance as well as radius variance was similar. For all accelerometer placements the mean radius was rarely used. This analysis also revealed that decision trees mostly used different spherical variance and variance of the radius near the tree roots. This points again to strong importance of variances. For between-subject level classifiers (see Fig.: 3.10), we observed strong dominance of spherical variances and variance of the radius. Our analysis also showed that resting and lower body groups are quite easily separated from other activities. On the other hand two groups of upper body activities are hard to differentiate which is confirmed by an increase in the tree complexity.

In future work we will proceed in two directions: (1) we will explore the influence of higher moments (e.g. skewness and kurtosis) on the classification accuracy of the models and (2) we will apply other interpretable classification methods on the extracted features.

## 5 Appendix

### 5.1 Procedure of spherical coordinates system selection

Let us introduce the following notation for a vector of acceleration represented in Cartesian coordinates associated with accelerometer:

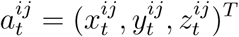

where: index *i* ∈ {1, …, 47} corresponds to a participants number, index *j* ∈ {1, 2, 3}correspond to a placement of an accelerometer (1 - hip, 2 - left wrist, 3 - right wrist), index *t* ∈ {1, 2, ⋯, *T*_*i*_} is the time index (*T*_*i*_ is the duration of an experiment for *i*-th participant). As was mentioned earlier the data from each accelerometer are measured in Cartesian system associated with device. In order to determine a spherical coordinate system, we rotated the Cartesian system associated with accelerometer so that the x, y and z axes coincide with (from participants point of view) forward, left and up directions, respectively (see Figure 2.2). The idea of such regularization largely borrows from Xiao et al. (2015). In second step we transformed the data to spherical coordinate system for which the zenith and azimuth directions corresponded to y and z axes of rotated Cartesian system, respectively. To establish the rotated Cartesian system we took few seconds of a signal for two activities: “standing still” and “lying still”. During this activities only the gravitational acceleration was detected by device. Therefore we assumed that the mean vector for activity “standing still” coincide with participants down direction and that the mean vector for second activity is close to backward direction. Let’s introduce the following notation for the mean vectors

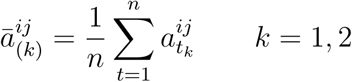

associated with “standing still” (1) and “lying still” (2) activities, respectively. Using above assumptions, we can set three basic directions (up, forward, left) according to the following equations:

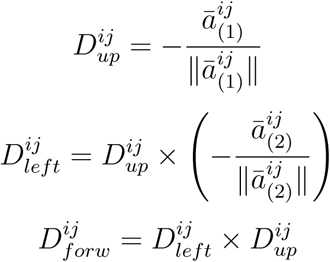

where *b × c* is a vector product of *b* and *c*. The relationship between rotated Cartesian system and associated with accelerometer has following form:

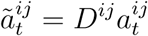

where 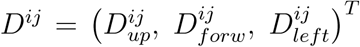 is the rotation matrix. After transformation of the data to a rotated Cartesian system we establish the spherical coordinates system according to the classical relationships:

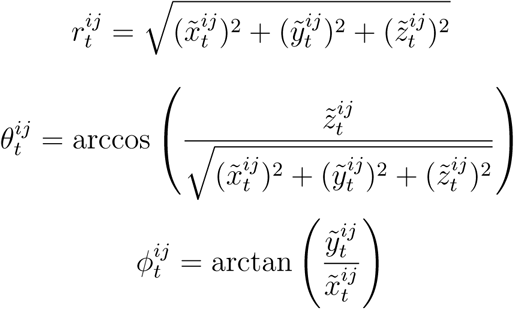

This procedure yields a representation of a vector of acceleration in a spherical system:

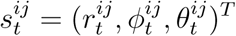

#### 5.2 Derivation of expression for the spherical variance and a proof of its rotational insusceptibility

Let’s recall the definition of the spherical variance:

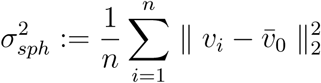

The following sequence of transformations proves presented in a paper expression for the spherical variance:

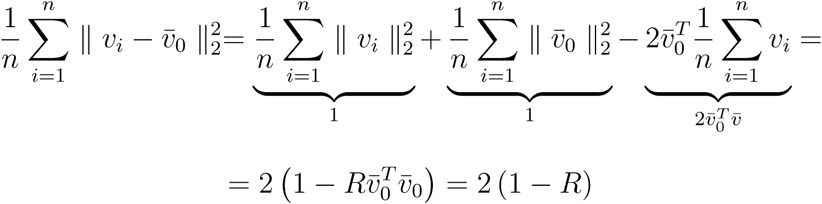

First equation is a consequence of bilinearity of the scalar product and its relationship with the norm. In second we use the assumption that the vectors 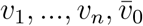 are normalized and a formula 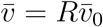 The last equation is again a consequence of a fact that 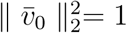 From above it is clear that in order to prove the rotational independence of the spherical variance it is sufficient to show rotational insusceptibility of *R*. Let’s recall that:

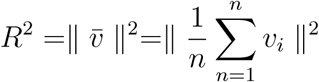

where *v*_*i*_ = (*x*_*i*_, *y*_*i*_, *z*_*i*_)*^T^* was a Cartesian representation of i-th point on a sphere. Let’s denote the same set of points in a fixed, arbitrarily chosen, rotated system in a following way 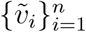. From linear algebra we know that there exist unique matrix of rotation *S* connecting points from both sets:

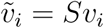

Now let’s consider the 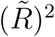 in rotated system:

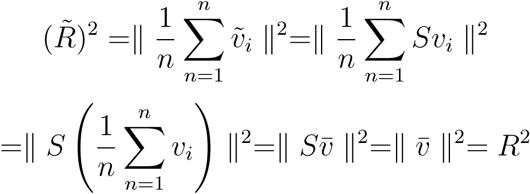

From above sequence of equations one can see that the choice of spherical coordinate system has no influence on the values of the spherical variance.

